# Effective Ensemble of Deep Neural Networks Predicts Neural Responses to Naturalistic Videos

**DOI:** 10.1101/2021.08.24.457581

**Authors:** Huzheng Yang, Shanghang Zhang, Yifan Wu, Yuanning Li, Shi Gu

**Affiliations:** Department of Computer Science and Engineering, University of Electronic Science and Technology of China, Chengdu, China; University of California, Berkeley, CA, USA; University of Pennsylvania, Philadelphia, PA, USA; Department of Neurological Surgery, Weill Institute for Neurosciences, University of California, San Francisco, CA, USA

**Author notes:** Correspondence to: Yuanning Li < >, Shi Gu < >.

## Abstract

This report provides a review of our submissions to the Algonauts Challenge 2021. In this challenge, neural responses in the visual cortex were recorded using functional neuroimaging when participants were watching naturalistic videos. The goal of the challenge is to develop voxel-wise encoding models which predict such neural signals based on the input videos. Here we built an ensemble of models that extract representations based on the input videos from 4 perspectives: image streams, motion, edges, and audio. We showed that adding new modules into the ensemble consistently improved our prediction performance. Our methods achieved state-of-the-art performance on both the mini track and the full track tasks.

## 1. Introduction

Building computational models for the human perception system is an important step towards the ultimate goal of better understanding human intelligence and guiding artificial intelligence engineering. Previous studies have shown that pre-trained task-optimized deep neural networks can extract useful representations which lead to neural encoding models of single static images with high performance (Yamins & DiCarlo, 2016; Schrimpf et al., 2020). However, the human visual system does not operate on independent static images but on continuous stimuli of videos. *The Algonauts Challenge 2021* (Cichy & Dwivedi, 2021) aims to improve our understanding of the neural basis and computational models of visual cognition, particularly regarding to naturalistic stimuli from everyday events. The goal of the challenge is to predict neural response to video clips of daily events recorded by functional magnetic resonance imaging (fMRI) at the voxel level. Motivated by the success of ensemble learning and multimodal methods (Zhou, 2019; Baltrušaitis et al., 2018), we develop an ensemble model that combines feature representations in deep neural networks from multiple perspectives and modalities, including image streams, motion, edges and audio features. We show that representations from each perspective separately improves the prediction performance of the ensemble encoding model. Our results also demonstrate that different brain areas integrate feature information on different spatial scales and show distinct levels of selectivity for these feature perspectives. The code of our implementation is available at https://github.com/huzeyann/huze_algonauts.

## 2. Dataset and Setup

In the challenge, the stimulus set includes a total of 1102 videos, 1000 for training and 102 held-out for online submission. These videos are 3-second clips of daily events (Cichy & Dwivedi, 2021). For the 1102 videos, most of them are recorded at 256×256 resolution and 30 fps for 3 seconds, with about 100 videos without audio and 200 videos blow 25 fps.

We trained our models on the first 900 video-fMRI pairs and used the rest 100 as validation set. The validated optimal model is used for the final online evaluation.

## 3. Methods and Experiments

We built an ensemble of backbone models from different feature perspectives (see Fig.1). Our final model with the best prediction performance consisted of 2 phases (see Fig.2):

**Figure 1.**
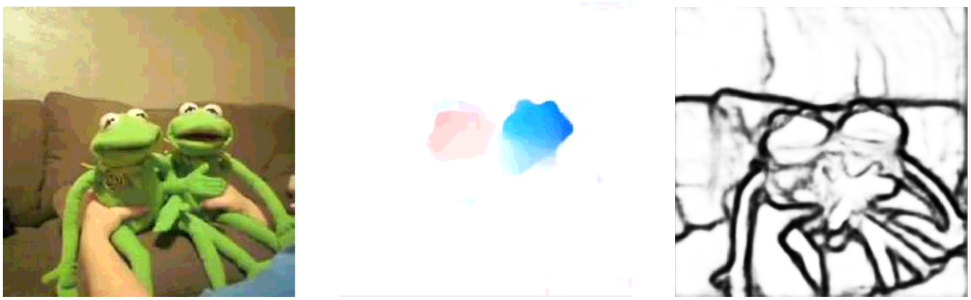
Examples of different perspectives of features. *(left)* Image features based on RGB frame; *(mid)* Motion features by optical flow estimation; *(right)* Edge features from perceptual edge detection.

**Figure 2.**
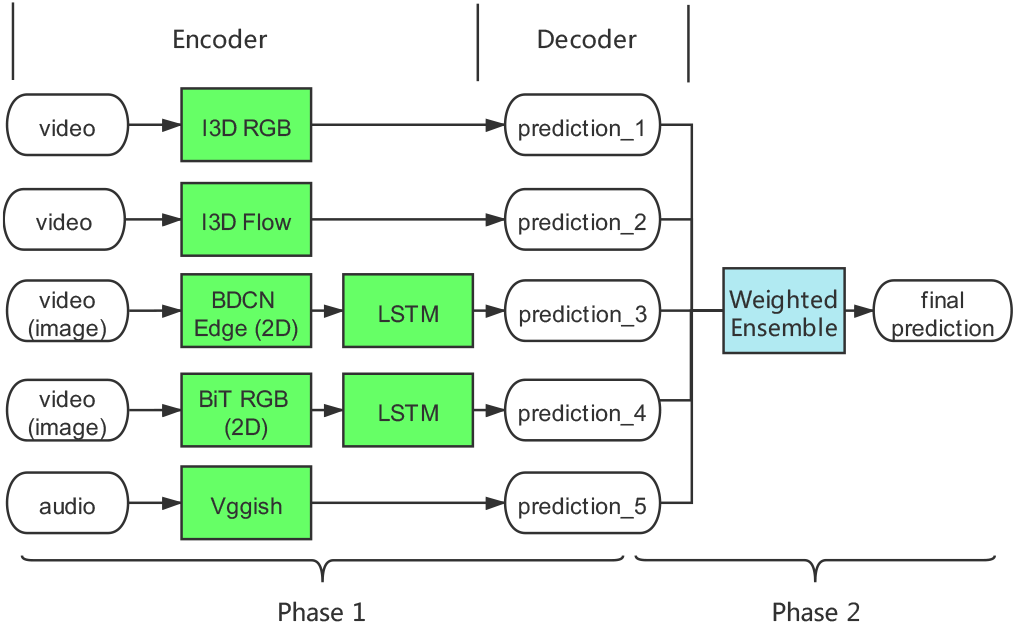
Schematics of the 2-phase ensemble model.

1. Training of encoding models on different perspectives.
2. Ensemble of predictions from all encoding models.

### 3.1. Ensemble Framework Overview

Our proposed model consists of a set of different feature encoder and decoder networks. These networks learn feature representations of the stimulus videos from different perspectives, including image streams, motion, edges, and audio. The feature encoders take in the stimulus video and extract feature vectors that contain relevant information. The decoders then take in these extracted feature vectors and generate predictions of neural responses.

To optimize our model performance, we performed two phases of model ensemble. In phase 1, we made the ensemble over different hyperparameters and architectures within the set of models from the same scope of feature modality. In phase 2, we took the ensemble models from phase 1 and combined them into the final neural encoding model.

The ensemble procedure assigned a weight to each model. The weights were optimized by differential evolution (Das & Suganthan, 2010) to maximize the correlation score on the validation set. This method of weight ensemble enables better performance on the validation set.

### 3.2. Encoder Models

We used 5 different types of encoder networks in our proposed framework: 1) Inflated 3D ConvNet image stream (**I3D RGB**): extracting the spatio-temporal information in the image stream; 2) Inflated 3D ConvNet optical flow stream (**I3D Flow**): extracting the motion dynamic information from the videos; 3) Bi-Directional Cascade Network for edge detection (**BDCN Edge**): extracting edge and contour information from the images; 4) Big Transfer (**BiT**): additional general image representations; 5) Convolutional network for audio classification (**Vggish Audio**): extracting audio information.

#### 3.2.1. I3D RGB

To extract the spatio-temporal features from the videos, we used the 3D Resnet model proposed in (Monfort et al., 2019). We fine-tuned the 3D Resnet model pre-trained on multi-label version of the full Moments in Time Dataset (Monfort et al., 2019). This model was pre-trained at 224×224 resolution and 16 frames, and we fine-tuned it for 288×288 resolution and 16 frames. We took the output of each of 4 Resnet blocks (res1 … res4) and connected the intermediate layers to an adaptive pooling layer with various pooling sizes, then flattened the output and fed it to the decoder model to make predictions. We tuned the backbone model end-to-end at half the learning rate after freezing it for 4 epochs.

##### Single-layer Encoder

The size of layer outputs from the Resnet blocks were res1: 256×8×72×72, res2: 512×4×36×36, res3: 1024×2×36×36, res4: 2048×1×9×9. We took each of these 4 outputs and employed adaptive pooling to reduce their temporal-spatial dimension to 1 × *k* × *k*. We ran a grid search on the pooling size *k* to find the proper pooling size for each layer and each ROI based on their validation correlation (Fig.3). For V1, V2, and V3, lower level features (res2, res3) and larger pooling size (smaller receptive field size) yielded better results, while EBA, LOC, PPA, FFA, and STS preferred higher-level features (res3, res4).

**Figure 3.**
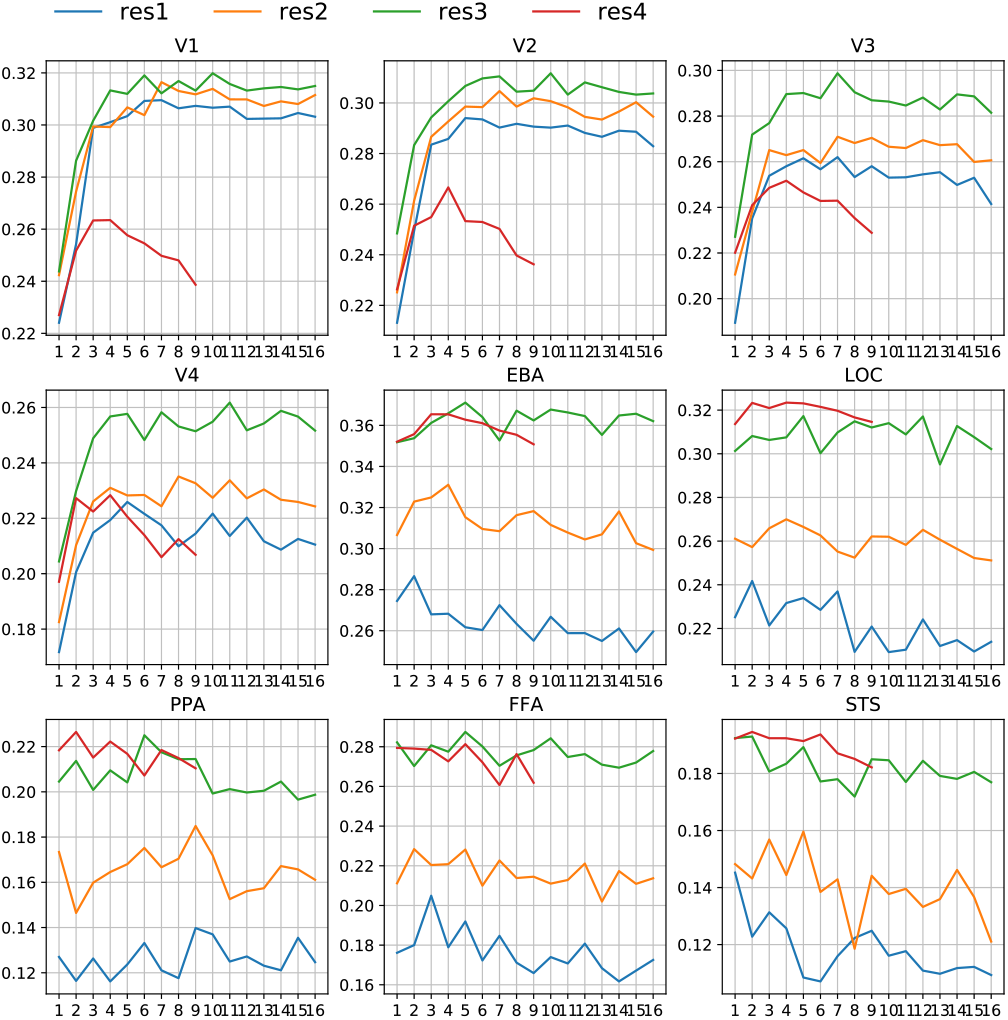
Summary of results using different pooling parameters in I3D RGB layers. Pooling size (x-axis) and layers (colored lines), validation correlation score (y-axis). A larger pooling size means a smaller receptive field size and more spatial information. The later layer represents higher-level features.

##### Multi-layer Encoder

We also built a multi-layer encoder through concatenating the outputs from different Resnet layers and employed the spatial pyramid pooling to summarize on multiple levels of features. The best multi-layer model slightly outperformed the best single-layer model only before the ensemble of different hyperparameters. After the ensemble of models with all searched hyperparameters, the performance of these 2 version of models on the validation set showed no significant difference (see Fig.4).

**Figure 4.**
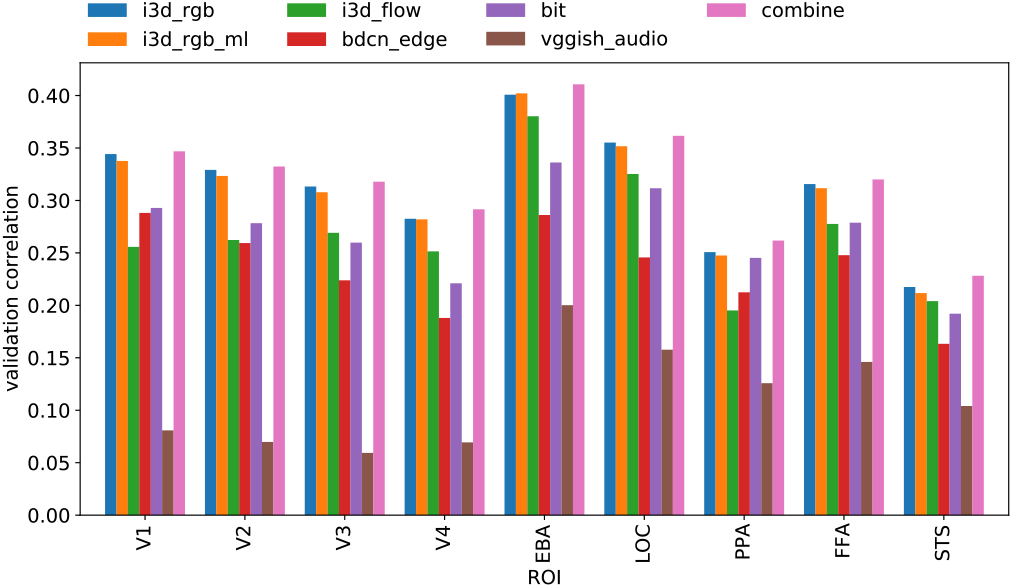
The performance of single models in each ROI. Different colors indicate different models. X-axis is ROI and y-axis is the correlation score on the validation set.

#### 3.2.2. I3D Flow

To include more dynamic information from the videos, we employed networks that extract information based on optical flows from the stimuli. Specifically, we took the flow stream of the 3D Resnet model pre-trained on Kinetics400 Dataset as proposed in (Carreira & Zisserman, 2017). The optical flow used as input was generated by RAFT (Teed & Deng, 2020). This model was pre-trained at 224×224 resolution and 64 frames, we fine-tuned at native 256×256 resolution and took the center 224×224 crop (we observed noise in edges when generating optical flow, center crop eliminated such noise and led to better performance). For videos that had less than 64 frames (videos recorded at 15fps), we employed **ffmpeg**’s temporal interpolation to up-sample them to 22fps, this up-sampling step led to more noise when generating optical flow. We did not tune this backbone model end-to-end due to implementation difficulties. The later part of the I3D Flow model was identical to the I3D RGB model: we ran a search on layers and pooling sizes, then ensembled all models in the search scope.

#### 3.2.3. BDCN Edge

Considering the sensitivity to edges in the early visual cortex and the object category selective properties in the ventral temporal areas, we also included networks that extract object shape information such as edges. Following (He et al., 2019), we took the edge detection output and employed adaptive pooling on it before feeding it into an LSTM layer to construct the edge-based feature representations. This model was pre-trained by 500×500 crops. We ran a grid search of input resolutions in [32, 48, 64, 96, 128], frames in [4, 10], pooling size in [8, 10, 12, 14, 16, 20, 24, 28, 32], and ensembled all the models in the search scope. The backbone was tuned end-to-end after freezing for the first 5 epochs and optimized at a 0.25 learning rate ratio.

#### 3.2.4. BiT

To gain additional feature representations from the images, we took the intermediate layers of Resnet from the pretrained BiT-M-R50×1 model (Kolesnikov et al., 2020) and fed it into a LSTM layer to make predictions. The fine-tuning was made at 224×224 resolution and 4 frames with pooling size in [1, 2, 3, 4, 5, 6, 7]. The backbone is tuned end-to-end after frozen for 4 epochs, then trained at a 1.0 learning rate ratio.

#### 3.2.5. Vggish Audio

Considering areas such as STS may encode multimodal information, we also included networks that represent audio features from the video. Although the actual audio was not presented to the subjects, additional information about the video may be represented in the audio and may correlate with the neural responses. Following (Hershey et al., 2017), we took the final embedding output from their model (dimension 3 × 128) as input. For videos that did not have audio, we left their input to zeros with the same shape. The backbone was not tuned end-to-end considering the simple fact that the audio was not played during the fMRI scan.

### 3.3. Decoder Models

The decoder model takes feature vector from the encoders for each video and predicts the corresponding neural responses in each voxel. For the *mini_track*, we employed a fully connected neural network to make predictions of all voxels in the same ROI for all subject. For the *full_track*, with the addition of spatial coordinates, we employed a convolution transpose model to make predictions of the whole brain. The output channel of the last convolution layer was set to match the number of subjects in order to predict all subjects’ neural responses simultaneously.

## 4. Results

### 4.1. Submission Scores in Mini- and Full-tracks

Table 1 lists the normalized correlation scores in the final online evaluations when adding models from different perspectives to the ensemble procedure. The performance score kept increasing when new perspectives were added. Due to the limited number of submissions during the challenge, we did not make a full ablation analysis for each individual module. We leave it to future work. It’s worth noting that that the submission score improved when adding the audio model to the ensemble, though the audio was not played during the fMRI scan.

**Table 1.**
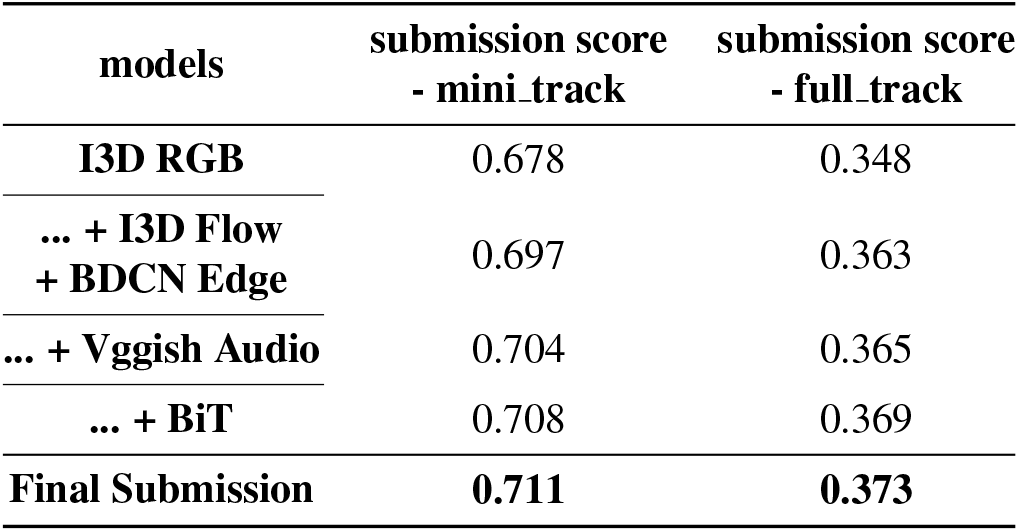
Summary of the submission scores using different ensemble strategy. The score kept improving when adding additional modalities. The final submission combined the best predictions for each ROI.

On average, our best ensemble models achieved normalized correlation score of 0.711 in the mini-track, and 0.373 in the full track. The best scores of ensemble model listed in Table 1 is slightly lower than our best submission scores because we combined the best predictions for each ROI when making the final submission.

### 4.2. Performance Scores of Inividual Models

In Fig.4 we plotted each model’s individual performance on the validation set for each ROI. I3D RGB was the best performing model, both single-layer and multi-layer versions of I3D RGB reached the best performance. Followed by BDCN Edge in earlier ROIs such as V1, V2, but this edge perceptual model had worse performance in higher areas, such as V3, V4, EBA, LOC, STS. I3D Flow had the best performance in STS, EBA, LOC, but performed worse than the edge perceptual model in V1. It’s worth noting that the audio model alone can make a moderate prediction for higher-level ROIs, such as PPA and STS, although audio was not played during fMRI scan. This suggests that there might be predictive coding of multimodal information in these areas, which is an open question to be studied.

### 4.3. Ensemble Weights of Individual Models

As described in 3.1, we used differential evolution to maximize the correlation score on the validation set to find the best ensemble weights. Fig.5 shows the final optimized ensemble weights, where V4, EBA, LOC, STS had a relatively high ensemble weight on Flow stream, and EBA, PPA, STS had a relatively high ensemble weight on the Audio stream. V1 had a higher ensemble weight on edge perceptual model. Note that i3d_rgb, i3d_rgb_ml, and BiT model may be overlapping with I3D RGB model regarding the extracted spatiotemporal features.

**Figure 5.**
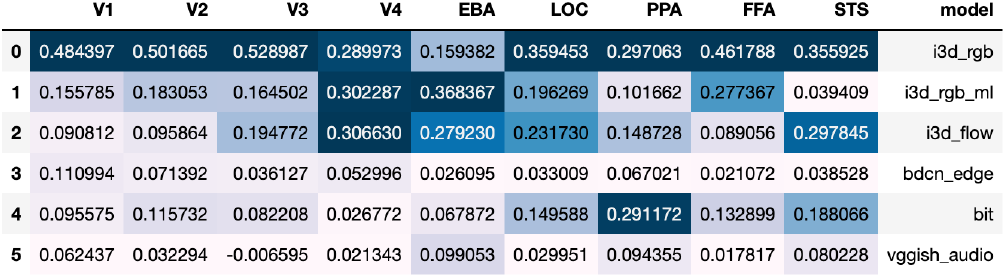
Ensemble weights for each model in each ROI. Each column sums to 1. Color is made column-wise, darker color means bigger value in ensemble weight.

### 4.4. ROI Results in the Full-track

During the challenge, since the mini_track voxels appeared in the full_track voxels as well, we retrieved their coordinates information via matching their fMRI responses. In practice, the replacement of the predicted values of the voxels in the full_track with their matched mini_track predictions led to a 0.015 improvement in submission score (from 0.306 to 0.321 in our earlier attempts).

We further compared scores within the same set of ROIs on the validation set for models trained on mini_track and full_track in Fig.6. It’s worth noting that even after the ensemble, the full_track model still performed worse compared mini_track model on the same ROI. This was also true when mixing several ROIs for training in mini_track, tested with I3D RGB model.

**Figure 6.**
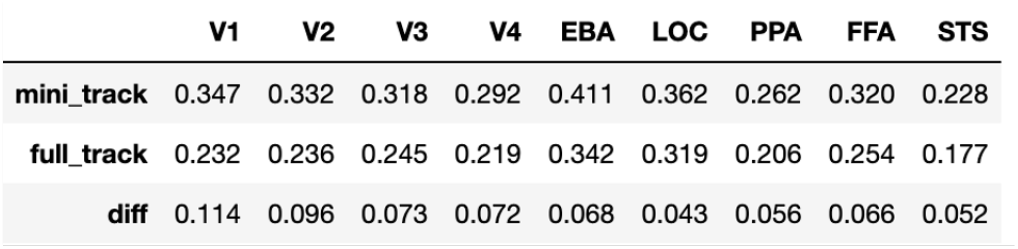
Comparison of the correlation scores of the same voxels between mini_track and full_track model, taken from best ensemble model evaluated on the validation set. full_track model perform significantly worse than mini_track model, differences are shown in the bottom row

Further experiments found that removing mini_track ROI voxels (around 30% of all voxels) during training for mini_track will lead to a significant drop in mean correlation score for the rest of voxels (0.11 drop to 0.09). This was true for the convolution transpose decoder model. The fully connected model, on the other hand, did not show the similar decline in performance. These accurate ROI voxels might give a boost to the other voxels during training through spatial smoothness or functional correlations in the activity.

We did replace the prediction of the ROI voxels in full_track with mini_track predictions in our final submission.

## 5. Implementation Details

AdaBelief Optimizer (Zhuang et al., 2020) was adopted to train our models, the base learning rate was set to 1e-4, beta=(0.9, 0.999), epsilon=1e-8, weight_decouple=True, weight_decay=1e-2 for non-bias weights. We set the batch size to 24 for mini_track and 32 for full_track. Mean-square error loss function was used to train encoder models, and early stopping conditioned on validation correlation. We used gradient accumulating trick to save memory if the model did not contain batch normalization or it was frozen. Pytorch’s native support for mixed-precision training (fp16) was employed for larger models (I3D RGB, I3D Flow, BiT). I3D RGB, I3D Flow, BiT models had 40M to 600M parameters depending on the pooling size. BDCN Edge model had less than 100M parameters due to the low dimensionality of the output edge prediction. Vggish Audio model had less than 10M parameters. The larger experiments were conducted on 1 NVIDIA RTX3090 GPU while smaller experiments are on 1 NVIDIA Titan Xp GPU, the largest model took 15 minutes to run 20 epochs before hitting early stopping.

## 6. Conclusion and Discussion

The key contribution of our proposed method is that we combined models from different modalities at the prediction level and improved the performance. We made the ensemble by directly optimizing the validation score with a global optimization algorithm. Fortunately, we did not observe severe overfitting on the validation set. Our results on the validation set and the final held-out testing were consistent except for STS and PPA, which were also the two ROIs with the lowest scores. However, our ensemble approach did not consider interactions between modalities at feature level, an integrated end-to-end model was harder to train and performed worse than the ensemble model in our experiments. Further work may consider training an integrated model that could outperform an ensemble of separately trained models.

## Notes

### Competing Interest Statement

The authors have declared no competing interest.

